# A partition-based spatial entropy for co-occurrence analysis with broad application

**DOI:** 10.64898/2026.02.23.707425

**Authors:** Adnane Nemri, Ovidiu Radulescu, Antoine Claessens, Thomas D. Otto

## Abstract

Despite the advent of spatial data science, including spatial biology, there exist few methods that study the distribution of points e.g. cells or individuals, accounting for both their own characteristics and environmental factors. We propose a new spatial entropy measure, termed the Regional Co-occurrence Entropy (RCE), that detects when categorical co-occurrences happen preferentially in specific environments. We demonstrate its use over a broad range of application fields. As examples, we study brain cell dynamics in Alzheimer’s Disease, identifying both known and likely novel interactions between immune cells around beta-amyloid plaques. We also investigate the diversity of buildings across a town neighborhoods, to detect potential drivers of social mixing at local scale. Finally, we dissect bird species distribution across a natural reserve, identifying potential vegetation-driven changes in community composition. Altogether, the proposed RCE enables rapid insights into interactions with an environmental component, making it a useful addition to the spatial data science toolbox.

**Significance Statement:** Spatial data is rapidly accumulating across fields as diverse as spatial biology, geography, ecology or astrophysics and promises to allow major scientific advances. For example, spatial biology is transforming our ability to study complex tissue organization and disease mechanisms. A key challenge remains the quantification of spatial relationships between individual points, such as cell types, in a statistically rigorous way. We propose a spatial entropy-based measure to quantify context-dependent interactions between categories, such as cell types across tissue subregions. Our novel method deriving from Information Science provides an efficient and versatile way to extract information from spatial datasets, with broad applicability across research fields.

## Introduction

As data capture technology progresses, we are witnessing increased generation and availability of spatial data across domains as diverse as satellite or drone imagery, ecological surveys or medical imaging (1). This is having a transformational impact on fields as diverse as health, law enforcement, warfare or environmental policy. A mixture of spatial statistics and artificial intelligence tools is typically used to automate the processing and analysis of such spatial data (2, 3). Advances in artificial intelligence offer powerful tools for tasks such as image pattern recognition and object counting. However AI remains computationally intensive, requires large training datasets and does not readily lend itself to hypothesis testing. Spatial statistics, on the other hand, can extract robust novel insights at low relative cost, allowing routine analysis of generated spatial data (4), such as live camera footage. Well-established fields such as single-cell omics have benefited from the recent addition of the spatial component (5). Until recently, most single-cell omics tools had to infer cell-to-cell interactions based on cell-type annotations and gene expression, without using spatial coordinates. With the added resolution gained from the cellular positions, one can now investigate how cells of the same type may behave differently depending on their tissue location. This has the potential to increase the signal-to-noise ratio and detect biologically meaningful spatial heterogeneity. Still, the integration of spatially resolved single-cell features and environmental information, for example tissue annotation, remains a challenge and an active field of research (6).

Among spatial statistics, spatial entropy (7) provides a simple way to incorporate spatial context and refine predictions of object-to-object interactions, with applications ranging from geography and ecology to molecular spatial biology (8). It builds on Shannon’s entropy (9), which measures the randomness in the occurrence of a phenomenon. While variants of Shannon’s entropy have been applied in computational biology (10), they remain insensitive to spatial organization. Spatial entropy overcomes this limitation by incorporating the arrangement of elements, allowing differentiation between aggregated patterns and uniformly random distributions (7).

To better understand spatially organized cellular interactions, we aimed to develop a spatial entropy measure capable of detecting whether specific pairs (or higher-order groups) of cell types co-occur more frequently in certain regions than others. Such a measure is critical for capturing region-specific interactions that may be overlooked by conventional analyses, providing deeper insight into tissue organization and cellular communication.

Existing spatial entropy measures capture spatial heterogeneity in occurrences of different types (categories) of objects (e.g. cell types), intensities of a phenomenon (e.g. immuno-fluorescence) or co-occurrences (7). Co-occurrences are defined as instances where pairs of objects of specific types (e.g. cell type A and cell type B) are found within a specified maximum distance. Batty’s entropy (11) uses partitioning of the observational plane as a factor, e.g. morphological annotation of tissues or landscape partitioning in GIS (Geographic Information System), to detect if a type of object is over/under-represented in some partitions, e.g. a cell type in sub-tissues within an organ. Yet, it is not suitable for co-occurrences of object type. When two types, A and B are both over-represented in partition *g_1_* compared to *g_2_*, it is not able to address whether individual points of A and B in partition *g_1_*are physically close (likely interacting) or scattered at random (independent recruitment). Conversely, Leibovici’s entropy (12) and Bayesian co-occurrence probability (13) capture co-occurrences of object types but ignore regional variations. Consequently, two nearby objects may be considered co-occurrent even when a physical barrier separates them, preventing their interaction.

To address this gap, we propose a spatial entropy measure for partition- and distance-based co-occurrences of object types. Unlike other tools that address the question of “who” in interactions, our tool also answers the questions of “where” and “how varied”. We demonstrate its utility across diverse spatial datasets, from microscale tissue architecture in medical applications to patterns of species interactions in spatial ecology and social diversity within human settlements. By providing more precise detection and spatial characterization of co-occurrences, the RCE approach has the potential to become invaluable across fields such as medical imaging, ecology, geomatics, climate modeling, agricultural landscape management, and the analysis of human mobility patterns. With its ability to model complex spatial relationships, this tool will drive new insights into tissue organization, species distribution, social dynamics, and environmental change.

## Results

### Definition of the Regional Co-occurrence Entropy (RCE)

We consider a two-dimensional space of total area *T*, divided in *G* partitions each of area *T_g_*, which may be disconnected, e.g. spots of sub-tissue *g_2_* inside a matrix of sub-tissue *g_1_* (see schematic representation in Fig. 1). *T\** is the area of the smallest partition. We consider *n* points (e.g. cells) forming a point pattern, with each point belonging to one of *I* possible categories (e.g. cell types) of a variable X. In our model, a co-occurrence is formed when *m* points physically locate at a distance less than *d* from each other within the same partition. As in Leibovici (12), we study a variable *Z* of co-occurrences of *m* points classified by their categories (e.g. for *m*=2, pairs of cell types *i_1_*and *i_2_*), where Z counts the co-occurrences *z_r_*. For *m*=2, the simplest case, *z_r_* represents pairs of X with *r* taking 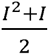 possible values. We consider only unordered co-occurrences, i.e. a co-occurrence *z_r_* of *i_1_* and *i_2_* is identical to a co-occurrence of *i_2_* and *i_1_* (to avoid the problem of double-counting). Note that the method is generalizable for *m*=3 (unordered triplets) and beyond. From Z, we derive a distance and partition-based co-occurrence variable *Z*|*L_Dg_*, where *L_Dg_* is a partition-based adjacency matrix of dimension *n* x *n*. We then calculate *p_zr_*_|*L*_*_Dg_* for each partition *g*, defined as the proportion of unordered m-tuples that occur in the same partition *g* and are separated by a distance less than *d*. The absolute entropy *H_RC_* of *Z*|*L_Dg_*, which we term the Regional Co-occurrence Entropy (RCE), is then calculated as

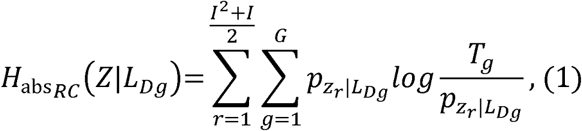

**Figure 1.**
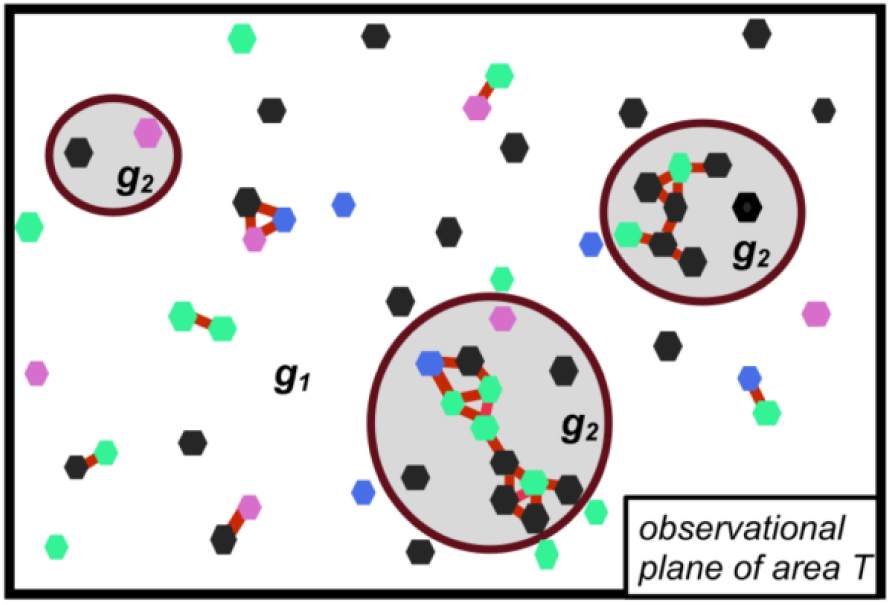
Schematical representation of partition and distance-based co-occurrences in an observational plane containing *G*=2 multi-polygon partitions (*g1* and *g2*). Co-occurrence pairs (*m*=2) at distance < *d* and in the same region *g* are shown using red links (-), for points with *I* = 4 categories (black, blue, purple and green hexagons).

and has the range 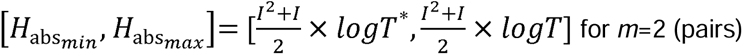 for *m*=2 (pairs). The relative RCE is calculated as

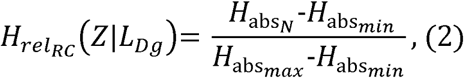

and has the range [0, 1]. Low RCE values mean specific pairs are over-represented in one or more partitions, whereas high values are obtained for uniformly distributed co-occurrences (relative entropy of ∼1), with near even densities across the differently sized partitions. After observing a lower than random RCE across the co-occurrence tuples, one would typically investigate which specific co-occurrence tuples contributed the most to lower the sum. Each of these 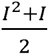 terms of the RCE sum are referred to as « decomposed » RCE for a specific tuple. The decomposed relative RCE’s vary in the range [0, 1] with the lowest decomposed RCE’s carrying the most signal. The relative RCE is thus the arithmetic mean of the decomposed relative RCE’s across tuples.

### Example application: cell dynamics in spatial biology

As a first example application, we investigated immune cells in Alzheimer’s disease. We used a *Xenium in situ* spatial transcriptomics model dataset of mouse brain (14), displaying extensive amyloid beta (*A*β) plaques, one of the hallmarks of the disease. As reported by the authors, astrocytes and microglia are detected in regions surrounding the amyloid plaques. From a morphology image, we identified the large compact *A*β plaques (red dots in Fig. 2a) and drew an expansion radius of 30 µm around them, which we termed “plaque regions”. The whole brain tissue slice (total of 810,334 µm2) was partitioned into a plaque (8% of total) and no-plaque (92%) map. After performing standard single-cell transcriptomics methods (see Methods), we focused our downstream analysis on 12 clusters (categories) of sub-types of astrocytes and microglia (7 and 5 clusters, respectively), totaling 10,335 cells (shown projected in Figure 2b,c). Based on the coordinates of cell centroids, 1,404 (13.5%) cells were in plaques and 8,931 (86.4%) outside of plaques. We calculated a relative RCE of 95.1% (lower means more signal) for the observed co-occurrences from the 12 categories across plaque and no-plaque partitions. To test whether this observed RCE was significant, we performed 1000 permutations of the cell categories (sub-types), while keeping their location fixed. The relative RCE for the simulations was significantly higher than for the actual distribution (mean of 98.6% with 1.3E^−4^% variance; Fig. 2d). Similarly, we found that the decomposed relative RCE per co-occurrence reached lower values for the actual distribution than for the simulations (minimum of 64.6% vs 84.6% in the simulations; Fig 2e).

**Figure 2.**
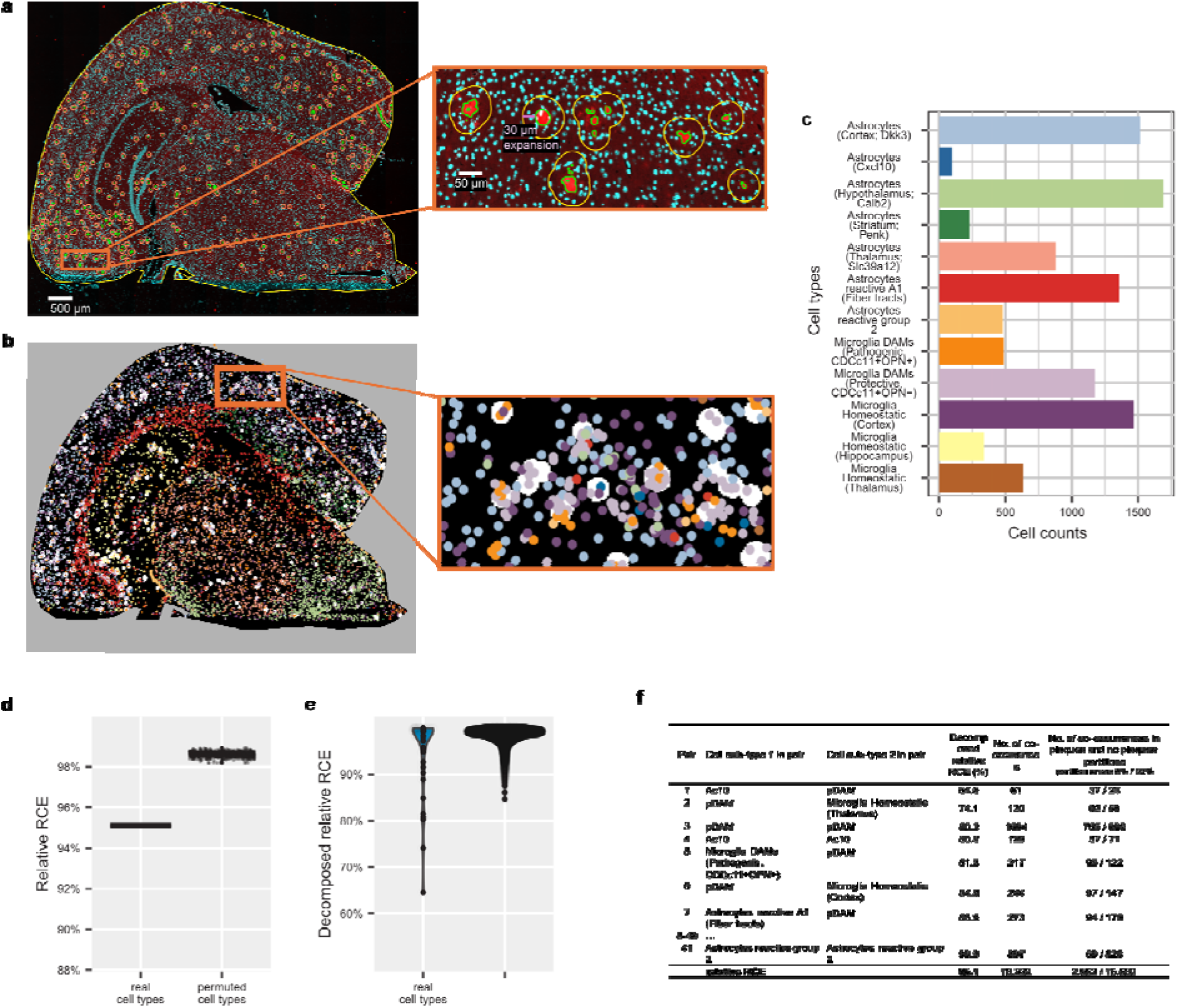
Migration dynamics of immune-related cell types in Alzheimer disease. (a) Immunofluorescent staining of diseased mouse brain, showing *A*β plaques (red), cell nuclei (DAPI, blue), 30 µm expansions forming the multi-polygonal “plaques” partition (orange); (b) projections of 12 selected immune-related cell sub-types, against plaques (white background) and no-plaques regions (black background) and (c) their cell counts; (d) relative RCE’s of the real cell type spatial distribution and 1000 random permutations of cell types (*d*=30, *min_coocs*=50); (e) distribution of decomposed RCE’s from each iteration; and (f) lowest decomposed RCE’s observed from the real cell type distribution.

We investigated further those specific co-occurrence pairs with lowest decomposed relative RCE (below 90%), all enriched in plaques and primarily involved the disease associated microglia sub-type “Protective DAMs (CD11c^+^OPN^−^)” (“pDAM”) and astrocytes sub-type “Astrocytes (Cxcl10)” (“Ac10”; Fig. 2f and Table S1). We divide them into homologous co-occurrence pairs (e.g. cell type A – cell type A) and heterologous pairs (e.g. cell type A - cell type B). The homologous co-occurrence pairs detected were pDAM-pDAM and Ac10-Ac10, indicating clumping of those cells in plaque regions. We counted 1694 pDAM-pDAM co-occurrences, incl. 765 in plaques (135 expected) and 929 outside of plaques (p-value untractable, ∼0). We found 128 Ac10-Ac10 co-occurrences, incl. 57 in plaques (10 expected) and 71 outside of plaques (p-value <10^−23^).

Notably, our findings suggest that pathogenic DAMs (CDCc11+OPN+) do not migrate to plaques to the same extent as pDAMs. pDAMs are known to gather next to large amyloid plaques (15, 16) and robustly ingest Aβ in a noninflammatory fashion (17). Microglia are also known to proliferate and activate one another via Csf1→ Csf1r signaling in the vicinity of plaques (18), suggesting this homologous pDAM-pDAM co-occurrence is biologically meaningful for cell-to-cell interactions.

Regarding astrocytes, we find that other astrocytes sub-types including the “reactive A1” and “reactive group 2” preferentially do not migrate to plaques. This could help resolve the current conflicting evidence about the migration of astrocytes towards plaques (19, 20). Enrichment of Ac10 cells in plaques is consistent with evidence that the chemokine CXCL10 is expressed primarily in astrocytes, localized around Aβ plaques in an AD mouse model (21) and, via its receptor CXCR3, drives neuronal pathogenesis (22).

The heterologous co-occurrence pDAM-Ac10 was also strongly enriched in “plaques”, suggesting no repulsion between those two cell types once they surround the plaques. This has biological relevance as microglia migration and activation has been proposed to be also driven by the signals they receive from neighboring astrocytes^12^, with which they form peri-plaque glial nets^13^. Taken together, our data suggest that, among astrocytes and microglia, specifically Ac10 and pDAM migrate to plaques, attracted by the plaque itself and by autocrine microglia signaling, where they interact to get activated and form peri-plaque glial nets. In addition, we observed the heterologous pairs pDAM – “Microglia Homeostatic (Thalamus)” and to a lesser extent pDAM – “Microglia Homeostatic (Cortex)” enriched in plaques regions (p-value <10^−27^ and p-value <10^−34^, respectively). This is suggestive of different dynamics of recruitment of homeostatic microglia towards activation into pDAM across the brain. In conclusion, the RCE and its decomposed terms have re-discovered previously known immune cell dynamics as well as uncovered potentially novel interactions. These findings help refine our understanding of differential cellular migration towards plaques and cell-to-cell interactions specific to the plaques environment, as well as potentially guiding future investigations.

### Example application: diversity in residential buildings as a proxy for social mixing in geography

Next, we applied the RCE approach to geographical data. We wanted to test the null hypothesis that direct neighbours across different parts of a town display uniform levels of social diversity at the local scale. Using aerial drone imaging data of Dennery Village (St. Lucia) in the Caribbean (Fig. 3a, left) , we chose rooftops as proxies for wealth, whereby intact and uniform roofing (referred hereafter as *intact*) indicates higher wealth whereas damaged or disparate roofing (hereafter *damaged*) indicates lower wealth. The aerial image was used to delineate buildings and classify them based on their rooftop material (Fig. 3a, right). We divided the village based on landscape and landmarks, namely rivers and canals. The expectation is that these factors are not drivers of social diversity and therefore partitions delimited this way should harbour fairly uniform levels of diversity at the local scale. Following the building delineation and rooftype classification, we counted 711 « healthy metal », 70 « concrete cement » and 9 « blue tarp » rooftype buildings, which were regrouped into the *intact* category . There were 401 « irregular metal » and 125 « incomplete » rooftype buildings, which were regrouped into the *damaged* category. After assigning the buildings to 6 partitions, there remained a total of 1312 buildings, comprising 790 « intact » and 522 « damaged » buildings (table S3). To detect direct neighbours, we set a distance threshold *d* of 15 meters between buildings. We counted 2330 co-occurrences across the 6 partitions (*min_coocs* = 10). In this example with I=2 categories, the possible pairs include *intact-intact*, *intact-damaged* and *damaged-damaged.* Our interest lies primarily in the heterologous pair *intact-damaged*, as a proxy for socially diverse direct neighbours.

**Figure 3.**
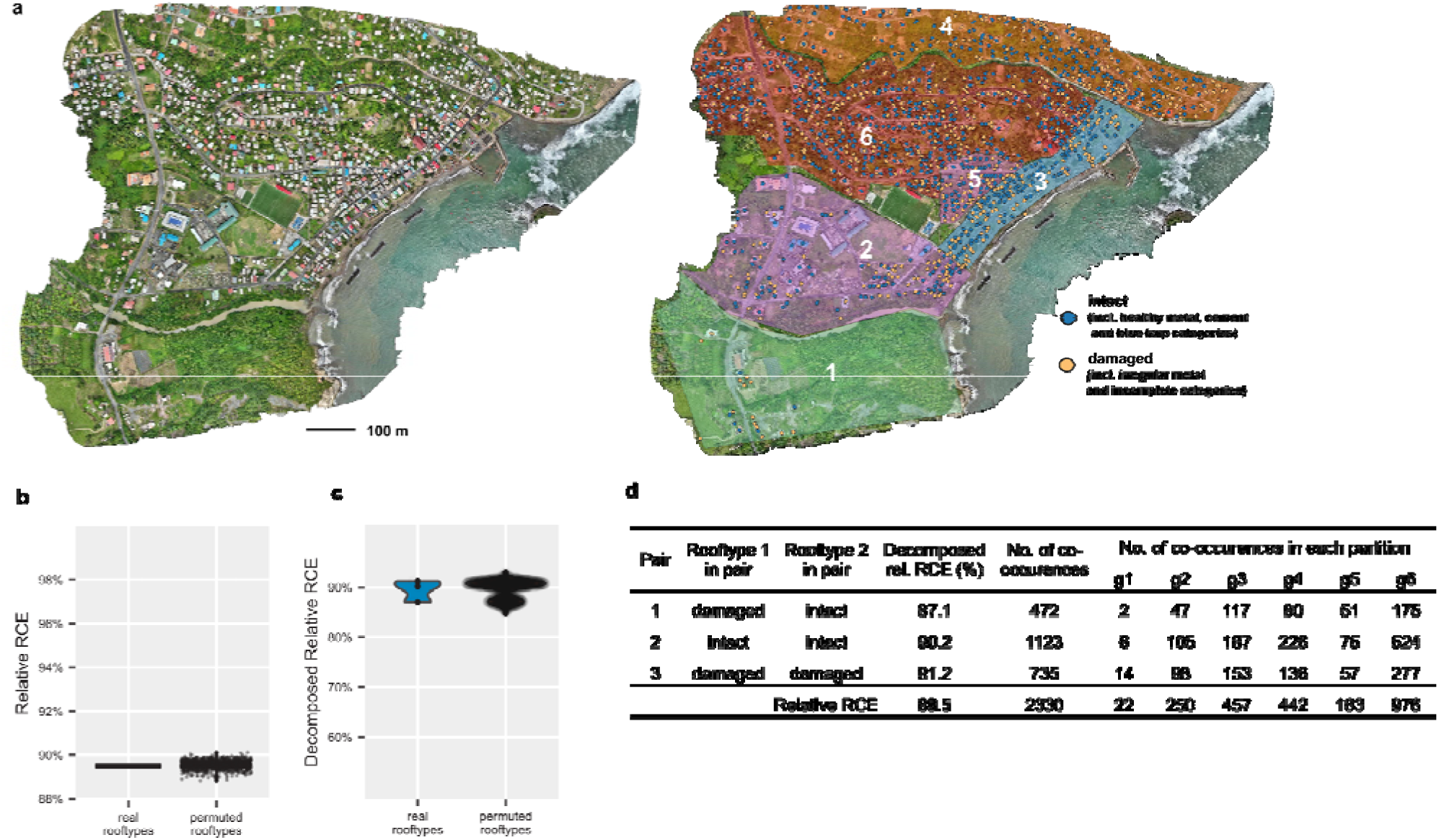
Rooftop diversity among direct neighbours compared across districts of Dennery Village (St Lucia). (a) drone image (left panel) and partitioning into districts and categorization of rooftops (right panel) (b) relative RCE’s of the real rooftype spatial distribution and 1000 random permutations of rooftypes (*d*=15, *min_coocs*=10); (c) distribution of decomposed RCE’s from each iteration; and (d) decomposed RCE’s observed from the real rooftype distribution.

The resulting relative RCE for the real rooftop distribution at 92 % was notably lower than in the Alzheimer example (95.1%). However it was firmly in the range of what could be observed in the random permutations (Fig. 3b), indicating that the frequencies of the 3 different co-occurrences did not vary significantly between partitions. Within partitions, we found that *intact-intact* and *damaged-damaged* co-occurrences were much more prevalent than intact-damaged co-occurrences. This clumping indicates that, irrespective of partitions, higher wealth households tend to have higher wealth neighbours and lower wealth households tend to have lower wealth neighbours. In the minority cases where higher wealth households have lower wealth neighbours, it occurs uniformly across the different town districts. As expected, our arbitrary partitioning of the town landscape, based on waterways, did not take into account the history of settlements and was found to not be a good factor driving social diversity. The null hypothesis of « no difference between partitions » is thus accepted. The lack of significant differences across partitions suggests that arbitrary geographic divisions (like rivers and canals) do not serve as important drivers of social diversity in this village.

Here, the RCE approach, especially applied as a comparison to that obtained from random permutations, therefore appears to be a rapid way to test hypotheses. One could envisage running sequentially the analysis using a series of partitioning methods to identify previously unknown factors driving diversity in human communities. This approach is also adaptable to different types of census data, making it a versatile tool for studying social dynamics in a spatial context.

### Example application: community composition in species ecology

Finally, we were interested to apply the RCE approach to ecological data, to investigate how the environment, in particular vegetation, drives bird community composition. We obtained observation data of 16 bird species on 90 sites located in the Disney Wilderness Preserve (Central Florida, USA; see Methods). We used the forest and grass cover to divide sites in 3 types. Here, the hypothesis was that some birds species pairs have partly overlapping habitat preferences and can be observed co-occuring in some environments. Similar to the Alzheimer example, the RCE for the actual distribution was significantly lower than for permuted data (Fig. 4d). The RCE observed for the real distribution was lower than that of 99.7-99.9% of random permutations, in two separate 1000 permutation tests, indicating structure in how bird communities are assembled. When looking at the decomposed RCE’s (Fig. 4e), less than 0.09% of 133,948 decomposed RCE’s from permuted bird species were below the lowest decomposed relative RCE observed for real data (60.6%). This indicates that both the RCE and decomposed RCE’s in this dataset are able to detect environment-driven species associations with certainty. We identified specific heterologous pairs of species that co-occurred preferentially in grass-dominant sites, e.g. Bachman’s Sparrow (BACS) - Common Ground Dove (COGD), or in mixed-dominance sites, e.g. Eastern Meadowlark (EAME) - Northern Mockingbird (NOMO; Fig. 4f, Table S2). As an example, neither BACS-BACS nor COGD-COGD are enriched in one specific environment (decomposed relative RCEs of 97.7% and 97.3%, respectively), however BACS-COGD are primarily seen in grass-dominant sites, suggesting that some specific factor in grass-dominant sites attracts both species. More generally, we can conclude that some environmental factor, likely ground cover vegetation, affects birds community composition and species interactions across the Wilderness Preserve.

**Figure 4.**
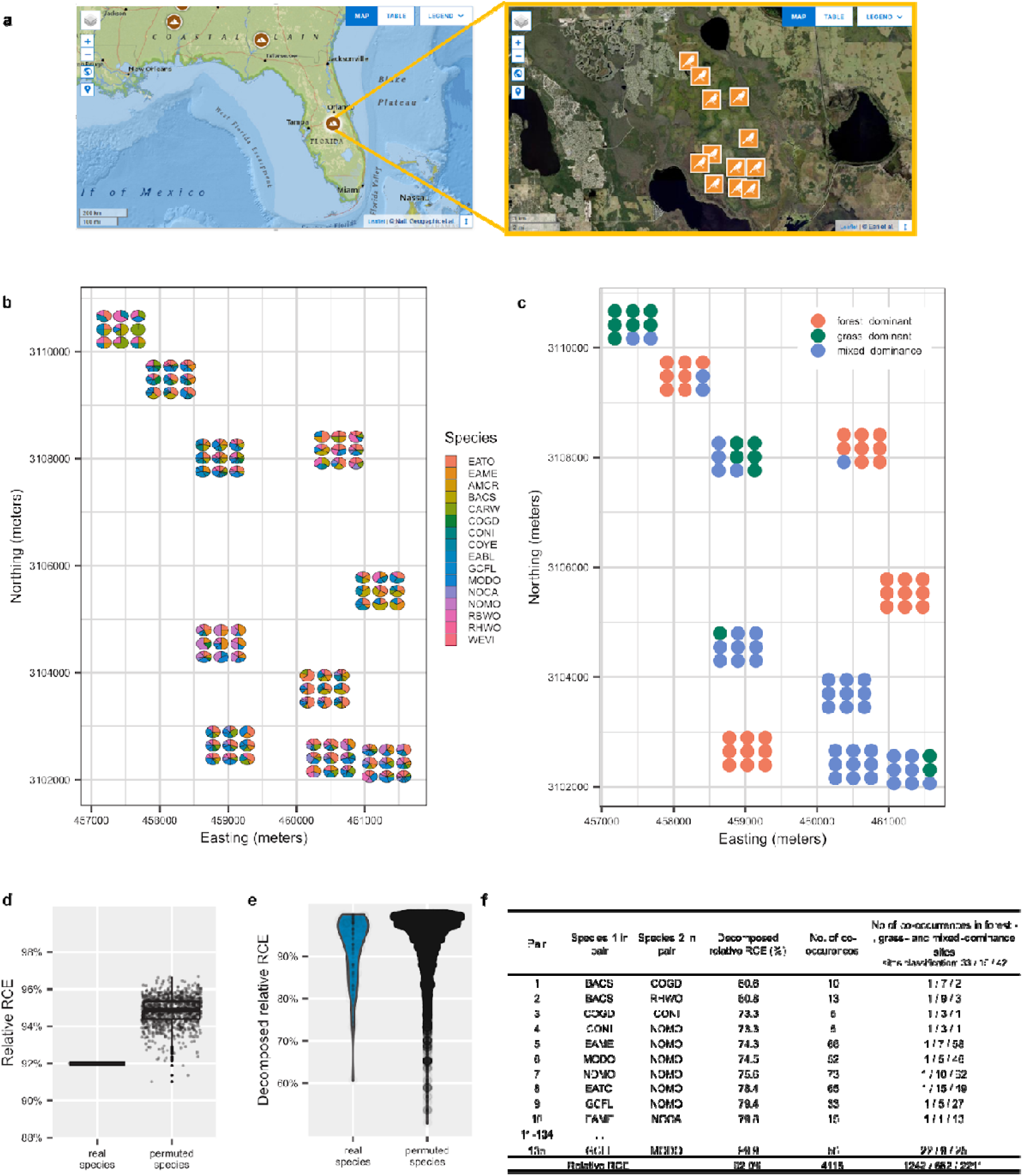
Bird community composition across a natural reserve. (a) Geographical location of 10 main sites in Disney Wilderness Preserve (Florida, USA), each sampled across 9 sub-sites; (b) bird communities composition and (c) environments classification (partitioning) of the sub-sites based on ground cover vegetation; (d) relative RCE’s of the real bird species distribution and 1000 random permutations of species (*d*=0, *min_coocs=*3); (d) distribution of decomposed RCE’s from each iteration; and (f) lowest decomposed RCE’s observed from the real species distribution.

## Discussion

Here, we introduce a spatial entropy framework that captures how co-occurrences of object types vary across regions, enabling the detection of spatially structured associations that remain inaccessible to existing approaches. By jointly incorporating partition structure and distance-based co-occurrence, the Regional Co-occurrence Entropy (RCE) quantifies where interactions occur and how consistently they are represented across environments, two dimensions of spatial organization that have traditionally been treated separately. This establishes a unified measure for investigating environment-dependent associations across diverse spatial systems.

The applications presented here illustrate the breadth and analytical power of this framework. In the bird ecology dataset, RCE highlighted specific species pairs, such as BACS–COGD, that consistently differed across habitats. Such region-specific pairings can motivate future ecological investigations into habitat-dependent behavioral or trophic interactions. In the Alzheimer’s tissue data, RCE recapitulated known plaque dynamics but additionally revealed spatially enriched co-localizations that had not been recognized previously, including pDAM microglia with Ac10 astrocytes. These findings suggest previously unexplored modes of microenvironment-specific cellular activation. Together, these results show that RCE can uncover fine-scale, spatially modulated interactions spanning cellular to ecological scales.

A notable feature of RCE is its generality and computational simplicity. The framework applies seamlessly from microscopic to geographic contexts and can naturally extend to 3D spatial data. Its ability to operate without specifying an interaction model stands in contrast to model-based approaches such as Gibbs point processes (23), which offer high precision but require extensive parametrization and computational cost. RCE therefore provides a complementary tool that enables rapid exploration of large spatial datasets, supports hypothesis generation, and facilitates the re-analysis of legacy datasets for overlooked spatial patterns.

Nonetheless, the RCE approach suffers from limitations, as discussed in detail in the Methods. There, we provide guidelines for the RCE parameters to ensure robustness and interpretability of the results as well as computational efficiency. In particular, the number of partitions G should be kept relatively low to prevent some partitions to harbor too few points. The threshold distance *d* should select only meaningful local interactions. The number of categories (*I*) considered and the number of categories in the tuple (*m*) can be relatively high, so long as the frequency threshold (*min_coocs*) is set conservatively. The aim is to prevent having to compute and interpret too high a number of tuples and also prevent very low-frequency tuples to have an outsized contribution to the overall RCE simply due to sampling effect. In any case, the use of random permutations of types allows to set a basis of expectation for the null hypothesis mirroring the specificities of the dataset at hand.

Overall, the RCE framework provides a versatile, domain-agnostic approach for quantifying environment-dependent associations across spatial datasets. By enabling the systematic comparison of co-occurrence patterns across regions, it offers a new analytic capability for uncovering structure and interactions in biological tissues, ecological communities, and human environments. As spatial data continue to expand in scale and diversity, RCE has the potential to become a foundational tool for analyzing the spatial organization of complex systems.

## Materials and Methods

### Algorithmic implementation

We implemented the method as a package in R (RC.entropy). To run, users provide a spatial point pattern of individuals (cells, buildings…) labelled as categories. They also provide a map indicating the partitions as well as a selection of the partitions of interest (*partition_sel ).* There are 3 user-chosen parameters input, which must be adapted to the data at hand.

1. *partitions_sel*, the list of *G* partitions selected to include in the analysis. After assigning individual points to their partitions, one may choose to exclude some partitions from the analysis. For example, we excluded the partition in our brain image which corresponds to an out-of-region area, shown as the light grey area around the brain in Fig. 1d (*partition_sel = [1,3]*). In the bird example, we segregated the 90 sites into three partitions and selected all of them (*partition_sel* = *[1,2,3]*). Generally, we advise against fragmentation of the observational plane into a large number of partitions, to improve the signal-to-noise ratio and the interpretability of the resulting entropy.
2. *d*, the maximum distance between two points to consider them co-occurrent. The distance *d* should be set to a meaningful value in the context of expected interactions between the points. When counts are recorded per site, i.e. individual points have the same coordinates, as in the bird example, this value *d* is set at 0.
3. *min_coocs*, the minimum counts of co-occurrences for a given categorical pair to include it in the analysis. As the Regional Co-occurrence Entropy formula is calculated using frequencies, it is sensitive to sample size, i.e. the number of co-occurrences observed for a given pair *r* across all partitions. Co-occurrences with very low counts tend to be skewed towards very low or very high relative entropies, stemming from sampling effect, despite low statistical robustness or information value. There are two cases where this occurs. First, the categories involved in the pair may have such low frequency that their co-occurrence has very low counts. Alternatively, regardless of their frequency, the categories involved in the pair may be very rarely seen in co-occurrences across all partitions, indicating either strong repulsion across all partitions or second-order environmental segregation, independent from the partitioning used presently. Our method is not able to detect such repulsion, although existing methods such as neighbourhood analysis may identify them. For second-order effects, another partitioning is required, based on a new hypothesis. In any case, we control for co-occurrences with very low counts across all partitions by applying a user-set threshold of minimum counts of co-occurrences for a pair (*min_coocs*). With increasing values of min_coocs, the low decomposed RCEs stemming for sampling effect are removed from the analysis, thereby removing noise and increasing signal (results not shown). To prevent a large number of low counts decomposed RCEs, we advise to keep the number of categories *I* low. Our method should hence perform better in terms of robustness and the total RCE across pairs will be more interpretable. This is to be balanced with the need for granularity in categories, for example by sub-typing cells in our first example, to allow novel insights.

Our current implementation is limited to *n*=46,340 points (e.g. cells or individuals), due to limits to storing *n*-by-*n* matrices. Possible improvements would include (1) storing the matrices differently to allow larger *n* values, (2) optimizing the time to compute the *n*-by-*n* pair-wise Euclidian distance calculations, which currently scales quadratically with *n*.

The current implementation runs on single-threaded CPU with minimal memory footprint for all three examples presented. The runtime to compute all 1000 permutations went from <5 minutes for the birds and village example to <2 hours for the Alzheimer’s example.

The RCE formula is sensitive to some edge cases, which were addressed in software engineering. Similar to other entropy methods, our equation is not able to handle cases where the frequency distribution includes 0, i.e. no counts are observed for one of the categories considered. In our case, this occurs when for a co-occurrence tuple, there are no counts in at least one partition. For example, in the case of a co-occurrence pair with *n*=100 counts where all 100 counts are seen in partition A and 0 in partition B, the calculation for partition B causes division by zero in the log(*T_B_*/p) term. If partition B is ignored from the summation, the decomposed RCE of log*T_A_*is obtained for partition A and for the RCE of the co-occurrence category across the partitions. Assuming A is the largest partition, log*T_A_*> log*T_B_*, with log*T_B_* being the minimal entropy that can be theoretically observed for these partitions. Arguably, a co-occurrence category with significant counts *n* that is never seen in one or more partitions, is very high signal and should instead have very low entropy, close or equal to log*T_B_*. Therefore, we chose not to exclude partition B from the summation. To this end, for such cases, we add 1 count to all partitions, e.g. the counts are now 101 in partition A and 1 in partition B, and the resulting entropy is ∼log*T_A_*, and tends closer towards it as *n* gets larger. If partition A is much larger than partition B (*T_A_* >> *T_B_*), then the RCE tends towards log*T_A_*∼log*T* and the relative RCE towards 1, as desired. Conversely, if all 100 counts are observed in the smaller partition B, and 0 in the larger partition A, using our transformation, the RCE tends towards log*T_B_* as desired (relative RCE of 0), indicating maximum signal.

Another edge case of our implementation occurs when the partitions are defined by multipolygons (connected or disconnected) and 2 points are within distance *d* from each other and in the same partition *g_i_*, however the gap between the 2 points is traversed by another partition *g_j_*. Although these points are within distance *d*, and are thus considered co-occurrent, they cannot physically interact, being separated by some kind of barrier between *g_i_* and *g_j_*. We do not currently control for this bias, which occurs rarely in our data and for our *d* values. One could control for it by annotating each polygon of a partition, *g_i,1_*, *g_i,2_,..,g_i,n_*, and only count as co-occurrences those that occur within the same *g_i,k_* sub-partition.

To estimate the probability that, for a given co-occurrence pair, the counts observed are expected at random, we calculated Poisson probability using the expected counts derived from relative area of the partition (using theoretical density as expected probability of event).

### Morphological annotation and alignment to single-cell in the mouse brain example

Immunofluorescent staining (IF) images and single-cell transcriptomics data were obtained from the 10X Genomics website (24). First, we loaded the IF image in QuPath v0.5.1 (25). We created a full image annotation. We then manually created a polygonal annotation around area of interest (“in_region”), and its inverse image (“not_in_region”; light grey area in Fig. 1d). A thresholder based on the red channel was created with a smoothing sigma of 5 and threshold of 1600 (Full resolution; Gaussian prefilter) and applied to classify pixels from the “in_region” layer and to find the red pixels groups corresponding to large compact *A*β plaques (minimum object size 2 µm^2^). The resulting “amyloid” annotation object was expanded 30 µm (constrained to parenting) to create the “expanded plaque” annotation. The inverse object “negative_expanded_plaques” was subsequently created. The “not_in_region”, “expanded_plaques” and “negative_expanded_plaques” were exported as geojson. They form the “not_in_region” partition (grey), “no plaques” partition (black) and “plaques” partition (white), shown in Fig. 1*d*, respectively. The geojson (“amyloid_expansion_30.geojson”) was imported using the spatstat package(26) as a tesselation and converted to a pixel image using a reduction factor of 16.

In parallel, we derived the cell locations, types and sub-types from the spatial omics data, following standard single-cell transcriptomics methods in Seurat(27). Briefly, we performed a first round of SCTransform, RunPCA (npcs=30), RunUMAP, FindNeighbors, FindClusters (resolution 0.3) on the 60,829 cells (Fig. S1). This resulted in 22 clusters, which we annotated manually using marker genes (using FindAllMarkers) (“mouse_brain_after_clustering.rds”). We then extracted the cells in the clusters 1 and 4, consisting primarily of astrocytes and microglia, to perform another round of clustering using SCTransform, RunPCA (npcs=30), RunUMAP, FindNeighbors, FindCLusters (resolution=1). Among the resulting 17 clusters, 14 clusters consisting of astrocytes sub-types and microglia sub-types were selected for downstream analysis (10,343 cells; Fig. S2). The remaining 3 clusters consisted of carry-over neurons and were removed from downstream analysis. From the Seurat object (“mouse_brain_after_2nd_clustering.rds”), we generated a spatial point pattern with cell sub-types used as marks.

Finally, the IF image was aligned with the single-cell image in Xenium Explorer following the recommended procedure (28) to obtain an alignment matrix as a csv file (“alignment_matrix.csv”). This alignment matrix contained the affine parameters (rotation, vertical and horizontal translations) and rescaling parameters, which, in combination with the pixel widths/heights of the IF image (0.325 µm) and single-cell image (0.2125 µm), were used to overlay the morphology pixel image and point pattern (cells). Cells located outside of the plaque and no plaque partitions, i.e. in the “not_in region” partition (grey area in 1D), were discarded. The final spatial point pattern consisted of 10,335 cells. The R workflow used to convert input data, count co-occurrences and estimate the RCE is available as the “alzheimer_example.Rmd” vignette.

### Human settlement example

A high resolution aerial drone image of Dennery Village (St Lucia) was obtained from Open aerial map (Dennery Village 3, ID: 67c21cadaaf1e28d21d2b778 (29)). From this image, buildings were transformed into a categorical spatial point pattern, while the landscape was transformed into polygons representing partitions.

Artificial intelligence models were used to identify buildings and classify them by rooftops, as per Tingzon et al (30). Specifically, building delineation was performed following the “Building Footprint Delineation for Disaster Risk Reduction and Response (Part 1)” tutorial (31) with minor modifications. A “output_polygons_raw.gpkg” file containing the polygons delimiting the identified buildings was obtained and minimally edited in QGIS v3.40.11 (32) to correct delineation errors, then saved as “output_polygons_edited_v1.gpkg”. Rooftop classification was performed following the “Rooftop Type and Roof Material Classification using Drone Imagery (Part 2)” tutorial (33). Centroids for the buildings were calculated using QGIS Geometry tools to obtain a “dennery_centroids.geojson” file.

The R workflow used to convert input data, count co-occurrences and estimate the RCE is available as the “village_example.Rmd” vignette.

### Bird example

The data were obtained from the spAbundance package (34). In short, 753 individual birds from 16 species were spotted across 90 sites from the Disney Wilderness Preserve (https://www.neonscience.org/field-sites/dsny), situated at the headwaters of the Everglades ecosystem in south-central Florida. The 16 species included in the dataset are: (1) EATO (Eastern Towhee); (2) EAME (Eastern Meadowlark); (3) AMCR (American Crow); (4) BACS (Bachman’s Sparrow); (5) CARW (Carolina Wren); (6) COGD (Common Ground Dove); (7) CONI (Common Nighthawk); (8) COYE (Common Yellowthroat); (9) EABL (Eastern Bluebird); (10) GCFL (Great-crested Flycatcher); (11) MODO (Mourning Dover); (12) NOCA (Northern Cardinal); (13) NOMO (Northern Mockingbird); (14) RBWO (Red-bellied Woodpecker); (15) RHWO (Red-headed Woodpecker); (16) WEVI (White-eyed Vireo). We allocated the 90 sites to 3 environments, using the covariate data of forest and grass coverage. Namely, a site was called “forest dominant” if its forest cover was 75% greater than its grass cover; “grass dominant’” if its grass cover was 75% greater than its forest cover, and “mixed dominance” otherwise. These 3 environments formed the partitions compared in this study. As a result, there were 33, 15 and 42 sites called “forest dominant”, “grass dominant”’ and “mixed dominance”, respectively. As the observations are summarized for a whole site, only coordinates of the center of the site are recorded. Therefore, we consider all individuals present on the same site to be co-occurrent. As a result, despite the relatively low total number of individuals (753), the number of co-occurrences is high and they represent a large diversity (135). For the data presented here, we set *min_coocs* = 3. The R workflow used to convert input data, count co-occurrences and estimate the RCE is available as the “birds_example.Rmd” vignette.

## Author Contributions

**Adnane Nemri**: Conceptualization, Software, Formal analysis, Writing - Original Draft. **Ovidiu Radulescu**: Formal analysis, Writing - Review & Editing. **Antoine Claessens**: Writing - Review & Editing, Supervision. **Thomas D. Otto:** Conceptualization, Writing - Review & Editing, Supervision, Funding acquisition.

## Classification

Physical and Biological Sciences; Applied Mathematics

## Data and code availability

The microscopy data, ecological data and drone images used in the article were all obtained from public datasets, as described in Methods. For the Alzheimer example, files incl. “mouse_brain_after_clustering.rds”,“mouse_brain_after_2nd_clustering.rds” from the single-cell analysis workflow were saved at https://zenodo.org/records/15966249 and https://github.com/sii-scRNA-Seq/RC.Entropy for reproducibility purposes.

Code to reproduce the experiments in this work is available under MIT Licence via GitHub at https://github.com/sii-scRNA-Seq/RC.Entropy and as vignettes in the *CRAN* R package RC.entropy.

## Acknowledgments

This work received support from the French government, managed by the Agence Nationale de la Recherche (ANR) as part of the investment programme “France 2030” under the reference “ANR-21-EXES-0005”, from the Occitanie Region, and from the ExposUM Institute of the University of Montpellier [AN & TDO]. This work also received support from the Wellcome Trust [grant number 104111/Z/14/ZR to TDO] and the Bernhard-Nocht Institute for Tropical Medicine.

## Supplemental Data

**Table S1.** Distribution of the decomposed RCE for the 41 pairs observed at a frequency higher than 50 counts (out a theoretical 78 possible pairs from 12 categories incl. self-pairs), ranked from lowest to highest.

| Pair | Cell sub-type 1 in pair | Cell sub-type 2 in pair | Decomposed relative RCE (%) | No. of co-occurrences | No. of co-occurrences in plaques and no plaques partitions<br>partition areas<br>64,663 / 745,761 |
| --- | --- | --- | --- | --- | --- |
| 1 | Astrocytes (Cxcl10) | Microglia DAMs (Protective, CDCc11+OPN-) | 64.6 | 61 | 37 / 24 |
| 2 | Microglia DAMs (Protective, CDCc11+OPN-) | Microglia Homeostatic (Thalamus) | 74.1 | 120 | 62 / 58 |
| 3 | Microglia DAMs (Protective, CDCc11+OPN-) | Microglia DAMs (Protective, CDCc11+OPN-) | 80.3 | 1694 | 765 / 929 |
| 4 | Astrocytes (Cxcl10) | Astrocytes (Cxcl10) | 80.8 | 128 | 57 / 71 |
| 5 | Microglia DAMs (Pathogenic, CDCc11+OPN+) | Microglia DAMs (Protective, CDCc11+OPN-) | 81.5 | 217 | 95 / 122 |
| 6 | Microglia DAMs (Protective, CDCc11+OPN-) | Microglia Homeostatic (Cortex) | 84.8 | 244 | 97 / 147 |
| 7 | Astrocytes reactive A1 (Fiber tracts) | Microglia DAMs (Protective, CDCc11+OPN-) | 88.9 | 273 | 94 / 179 |
| 8 | Astrocytes (Cortex; Dkk3) | Microglia DAMs (Protective, CDCc11+OPN-) | 90.3 | 318 | 103 / 215 |
| 9 | Microglia DAMs (Protective, CDCc11+OPN-) | Microglia Homeostatic (Hippocampus) | 91.5 | 59 | 18 / 41 |
| 10 | Astrocytes reactive group 2 | Microglia DAMs (Protective, CDCc11+OPN-) | 92.7 | 108 | 31 / 77 |
| 11 | Microglia DAMs (Pathogenic, CDCc11+OPN+) | Microglia Homeostatic (Cortex) | 94.8 | 88 | 22 / 66 |
| 12 | Astrocytes (Hypothalamus; Calb2) | Microglia DAMs (Protective, CDCc11+OPN-) | 95.5 | 106 | 25 / 81 |
| 13 | Microglia DAMs (Pathogenic, CDCc11+OPN+) | Microglia DAMs (Pathogenic, CDCc11+OPN+) | 95.8 | 629 | 144 / 485 |
| 14 | Astrocytes reactive A1 (Fiber tracts) | Microglia DAMs (Pathogenic, CDCc11+OPN+) | 97.2 | 80 | 16 / 64 |
| 15 | Astrocytes (Cortex; Dkk3) | Microglia DAMs (Pathogenic, CDCc11+OPN+) | 97.4 | 98 | 19 / 79 |
| 16 | Astrocytes (Hypothalamus; Calb2) | Astrocytes (Thalamus; Slc39a12) | 98.1 | 163 | 2 / 161 |
| 17 | Microglia Homeostatic (Hippocampus) | Microglia Homeostatic (Hippocampus) | 98.4 | 398 | 66 / 332 |
| 18 | Astrocytes (Thalamus; Slc39a12) | Microglia Homeostatic (Cortex) | 99.1 | 68 | 2 / 66 |
| 19 | Astrocytes (Thalamus; Slc39a12) | Astrocytes (Thalamus; Slc39a12) | 99.2 | 1061 | 33 / 1,028 |
| 20 | Astrocytes (Cortex; Dkk3) | Astrocytes reactive A1 (Fiber tracts) | 99.3 | 118 | 4 / 114 |
| 21 | Astrocytes (Hypothalamus; Calb2) | Microglia Homeostatic (Thalamus) | 99.3 | 57 | 2 / 55 |
| 22 | Astrocytes (Hypothalamus; Calb2) | Astrocytes reactive A1 (Fiber tracts) | 99.4 | 108 | 4 / 104 |
| 23 | Astrocytes (Cortex; Dkk3) | Astrocytes reactive group 2 | 99.4 | 77 | 10 / 67 |
| 24 | Astrocytes (Striatum; Penk) | Astrocytes (Striatum; Penk) | 99.4 | 261 | 10 / 251 |
| 25 | Microglia Homeostatic (Thalamus) | Microglia Homeostatic (Thalamus) | 99.5 | 684 | 87 / 597 |
| 26 | Astrocytes reactive A1 (Fiber tracts) | Microglia Homeostatic (Thalamus) | 99.6 | 224 | 27 / 197 |
| 27 | Astrocytes (Hypothalamus; Calb2) | Astrocytes (Hypothalamus; Calb2) | 99.7 | 2638 | 126 / 2,512 |
| 28 | Astrocytes reactive A1 (Fiber tracts) | Astrocytes reactive group 2 | 99.7 | 123 | 14 / 109 |
| 29 | Astrocytes (Hypothalamus; Calb2) | Astrocytes reactive group 2 | 99.8 | 99 | 11 / 88 |
| 30 | Astrocytes (Hypothalamus; Calb2) | Astrocytes (Striatum; Penk) | 99.8 | 91 | 5 / 86 |
| 31 | Astrocytes (Cortex; Dkk3) | Astrocytes (Hypothalamus; Calb2) | 99.9 | 122 | 7 / 115 |
| 32 | Astrocytes reactive group 2 | Microglia Homeostatic (Cortex) | 99.9 | 68 | 7 / 61 |
| 33 | Astrocytes reactive A1 (Fiber tracts) | Microglia Homeostatic (Cortex) | 99.9 | 123 | 8 / 115 |
| 34 | Microglia Homeostatic (Cortex) | Microglia Homeostatic (Cortex) | 99.9 | 1666 | 156 / 1,510 |
| 35 | Astrocytes (Hypothalamus; Calb2) | Microglia Homeostatic (Cortex) | 99.9 | 495 | 33 / 462 |
| 36 | Astrocytes (Thalamus; Slc39a12) | Microglia Homeostatic (Thalamus) | 99.9 | 113 | 8 / 105 |
| 37 | Astrocytes reactive A1 (Fiber tracts) | Astrocytes reactive A1 (Fiber tracts) | 99.9 | 2028 | 179 / 1,849 |
| 38 | Astrocytes (Cortex; Dkk3) | Astrocytes (Cortex; Dkk3) | 99.9 | 1826 | 158 / 1,668 |
| 39 | Astrocytes (Striatum; Penk) | Microglia Homeostatic (Cortex) | 99.9 | 59 | 5 / 54 |
| 40 | Astrocytes (Cortex; Dkk3) | Microglia Homeostatic (Cortex) | 99.9 | 442 | 34 / 408 |
| 41 | Astrocytes reactive group 2 | Astrocytes reactive group 2 | 99.9 | 897 | 69 / 828 |
| relative RCE |  |  | 95.1 | 18,232 | 2,652 / 15,580 |

**Table S2.**
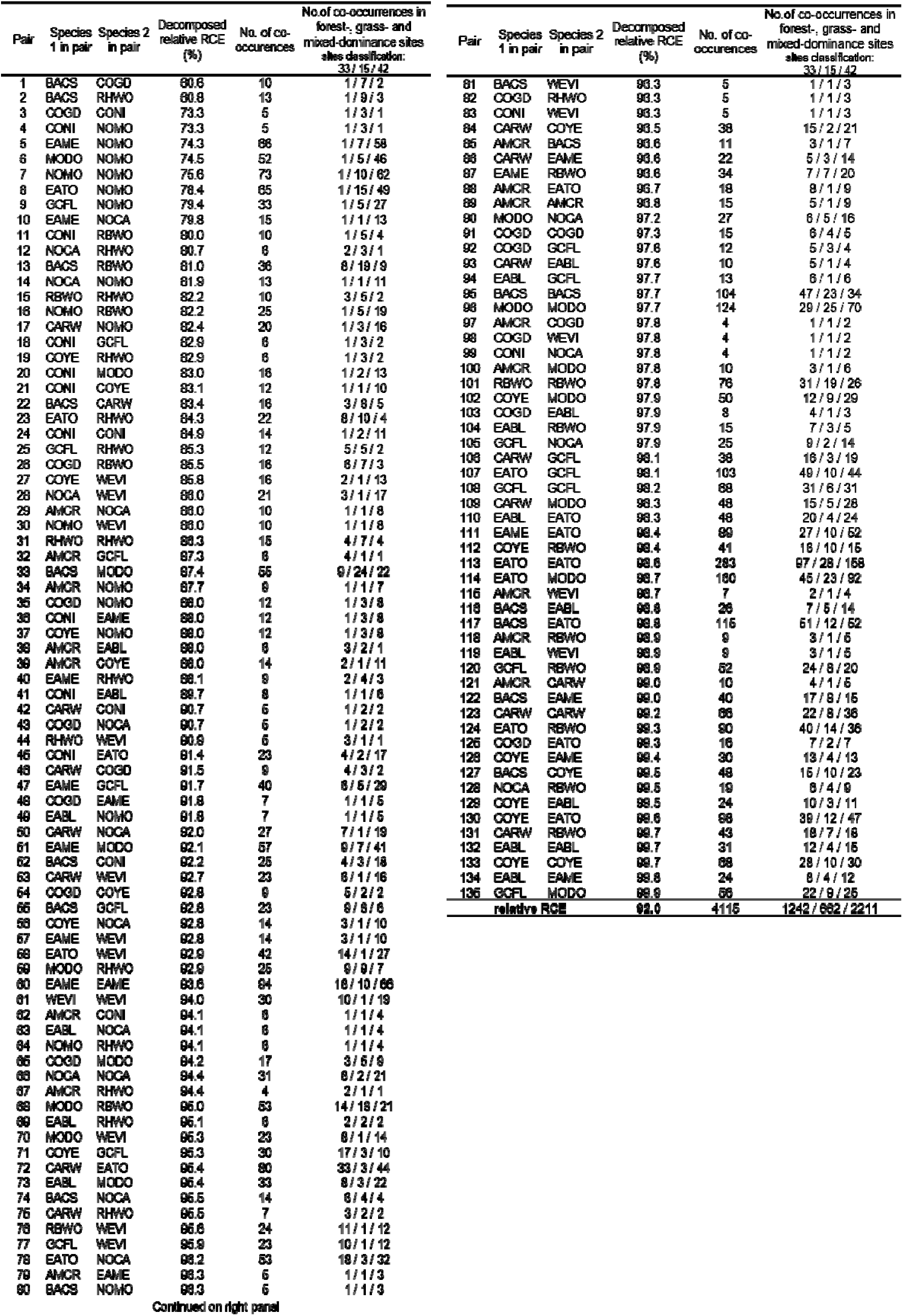
Distribution of the decomposed RCE for the 41 pairs observed at a frequency higher than 3 counts (out a theoretical 136 possible pairs from 16 categories incl. self-pairs), ranked from lowest to highest.

**Table S3.** Distribution of rooftypes across the 6 arbitrary partitions of Dennery Village (St. Lucia)

| Partition | No. of buildings | damaged | intact |
| --- | --- | --- | --- |
| 1 | 19 | 13 | 6 |
| 2 | 148 | 69 | 79 |
| 3 | 199 | 90 | 109 |
| 4 | 274 | 100 | 174 |
| 5 | 86 | 39 | 47 |
| 6 | 586 | 211 | 375 |
| Sub-total |  | 522 | 790 |
| Total |  |  | 1312 |

**Figure S1.**
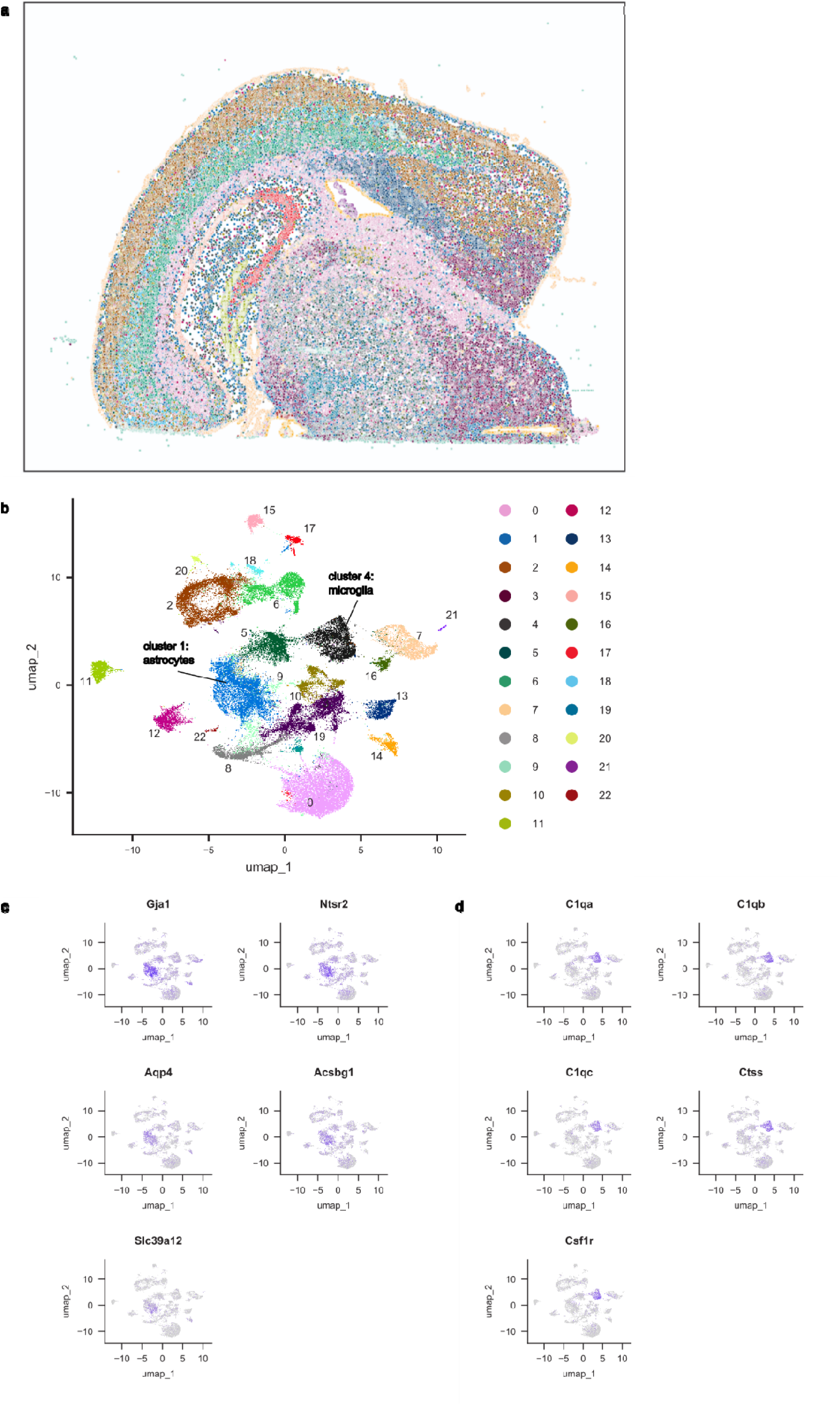
Cell types of a mouse brain with Alzheimer disease. Cells were projected using spatial coordinates (a), clustered based on transcriptomic profiling and manually annotated using marker genes (b). In particular, clusters specific for astrocytes (c) and microglia (d) were identified and subsequently subset for downstream analysis.

**Figure S2.**
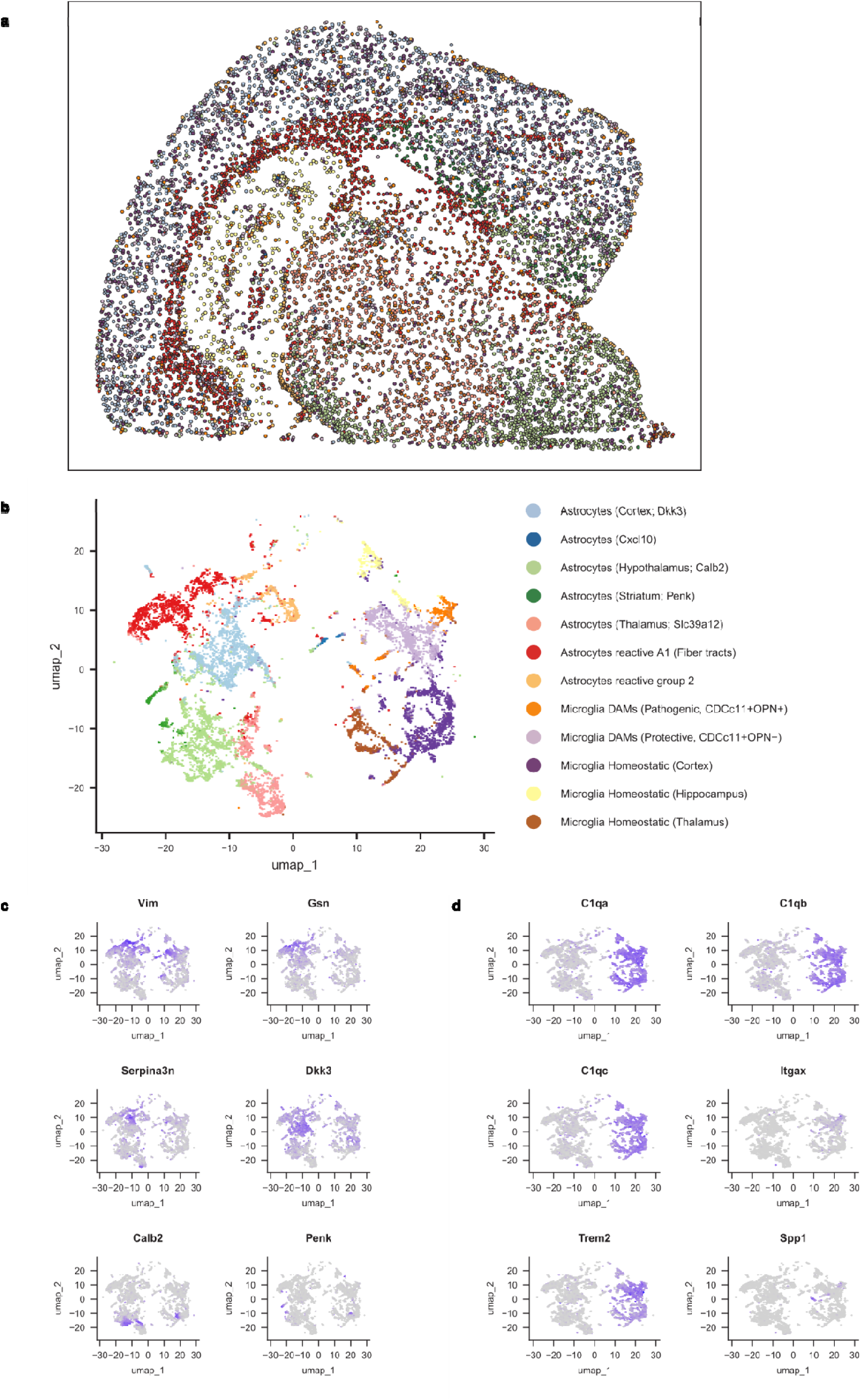
Microglia and astrocytes sub-types in a mouse brain with Alzheimer disease. Cells were projected using spatial coordinates (a), clustered based on transcriptomic profiling and manually annotated using marker genes (b), specific for astrocytes (c) and microglia (d).

**Figure S3.**
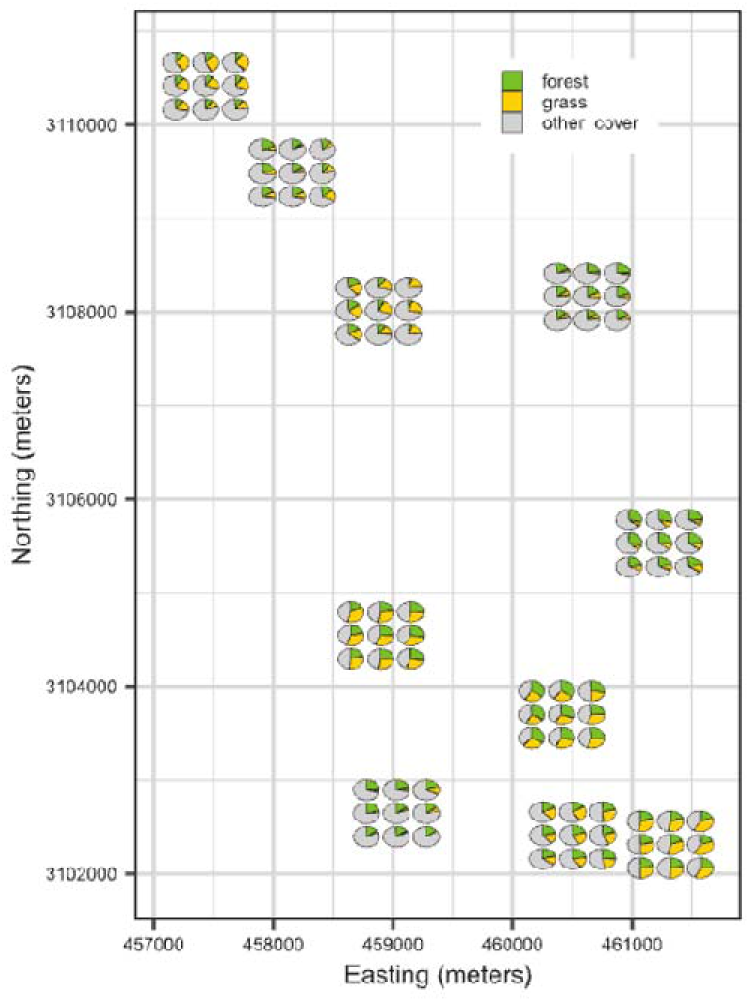
Ground cover vegetation ratios of sites across the Disney Wilderness Preserve (Florida, USA)

